# Prostate cancer peripheral blood NK cells show enhanced CD9, CD49a, CXCR4, CXCL8, MMP-9 production, and secrete monocyte-recruiting and polarizing factors

**DOI:** 10.1101/2020.07.06.189449

**Authors:** Denisa Baci, Matteo Gallazzi, Mortara Lorenzo, Annalisa Bosi, Giuseppe Buono, Angelo Naselli, Andrea Guarneri, Federico Dehò, Paolo Capogrosso, Adriana Albini, Douglas M. Noonan, Antonino Bruno

## Abstract

**Background:** Natural killer (NK) cells are effector lymphocytes of the innate immunity. Two major NK cell subsets are mostly present in the peripheral blood (pNKs): the cytotoxic CD56^dim^CD16^+^ NK cell subset (90-95% of pNKs), and the low cytotoxic, highly cytokine-producing CD56^bright^CD16^-/low^ NK cell subset (5-10% of pNKs). It has been demonstrated that NK cells in peripheral blood of patients with several tumors are altered. We have shown that in NSCLC and colon cancer, tumor associated circulating NK (pTA-NK) and tumor infiltrating NK (TI-NK) are skewed towards the CD56^bright^CD16^-/low^ phenotype. We have detected the production of pro-inflammatory and pro-angiogenic cytokines and chemokines. Other groups are reporting similar observations. There is still a lack of knowledge concerning the phenotype of pNK cells in prostate cancer (PCa). Here, we phenotypically and functionally characterized peripheral blood NK (pNK) from PCa patients (PCa pTA-NKs) and investigated their production of soluble factors, with endothelial cells and macrophage stimulatory action.

**Methods:** NK cell subset distribution was investigated in the peripheral blood of PCa patients, by multicolor flow cytometry (FC) for surface antigens expression. Protein arrays were performed to characterize the secretome on FACS-sorted pNK cells. Secreted products from FACS-sorted PCa TA-NKs were used to characterize their production of pro-inflammatory molecules. Secreted products from FACS-sorted PCa pTA-NKs were also used to stimulate endothelial cells and monocytes and macrophages, determining their ability to recruit and polarize them. Alterations of endothelial cells and monocytes, following exposure to secreted products from FACS-sorted PCa pTA-NKs, was assessed by RT-PCR. To confirm these observations, secreted products from 3 different PCa (PC-3, DU-145, LNCaP) cell lines were used to assess their effects on human NK cell polarization, by multicolor flow cytometry.

**Results:** Circulating NK cells from prostate cancer patients have been studied before, mostly for their impaired lytic functions. However, here we are the first to report that circulating pNK cells from PCa patients acquire a CD56^bright^CD9^+^CD49a^+^CXCR4^+^ phenotype with pro-inflammatory properties. We observed a similar polarization of heathy-donor derived pNK cells exposed to secreted products of three different PCa cell lines. Increased production of CXCL8, CXCR4, MMP-9, pro-inflammatory and reduced production of TNFα, IFNγ and Granzyme-B was detected. PCa TA-NKs released factors able to support angiogenesis *in vitro* and increased the expression of *CXCL8, ICAM-1* and *VCAM-1* mRNA in endothelial cells, confirming a pro-inflammatory signature. Secretome analysis revealed the ability of PCa pTA-NKs to release pro-angiogenic cytokines/chemokines involved in monocyte recruitment and M2-like polarization. In experimental setting, secreted products from PCa pTA-NKs can recruit THP-1 monocyte and polarize THP-1-differentiated macrophage towards *CD206/Arginase1/IL-10/CXCL8*-expressing M2-like/TAMs.

**Conclusions:** Our results show that PCa pTA-NKs are effector cells able to produce pro-inflammatory angiogenesis factors able to stimulate endothelial cells, attract monocytes and polarize macrophage to an M2-like type. Our data provides a rationale for the possible use of pNK profiling in clinical studies on PCa

**GRAPHICAL ABSTRACT:** Graphical Abstract:
Representative cartoon illustrating the pro-angiogenic features of PCa pTA-NKs.
A) direct effects of PCa pTA-NKs in supporting angiogenesis by interacting with endothelial cells. B) Proposed model for PCa pTA-NK pro-angiogenic activities via macrophage recruitment and polarization.

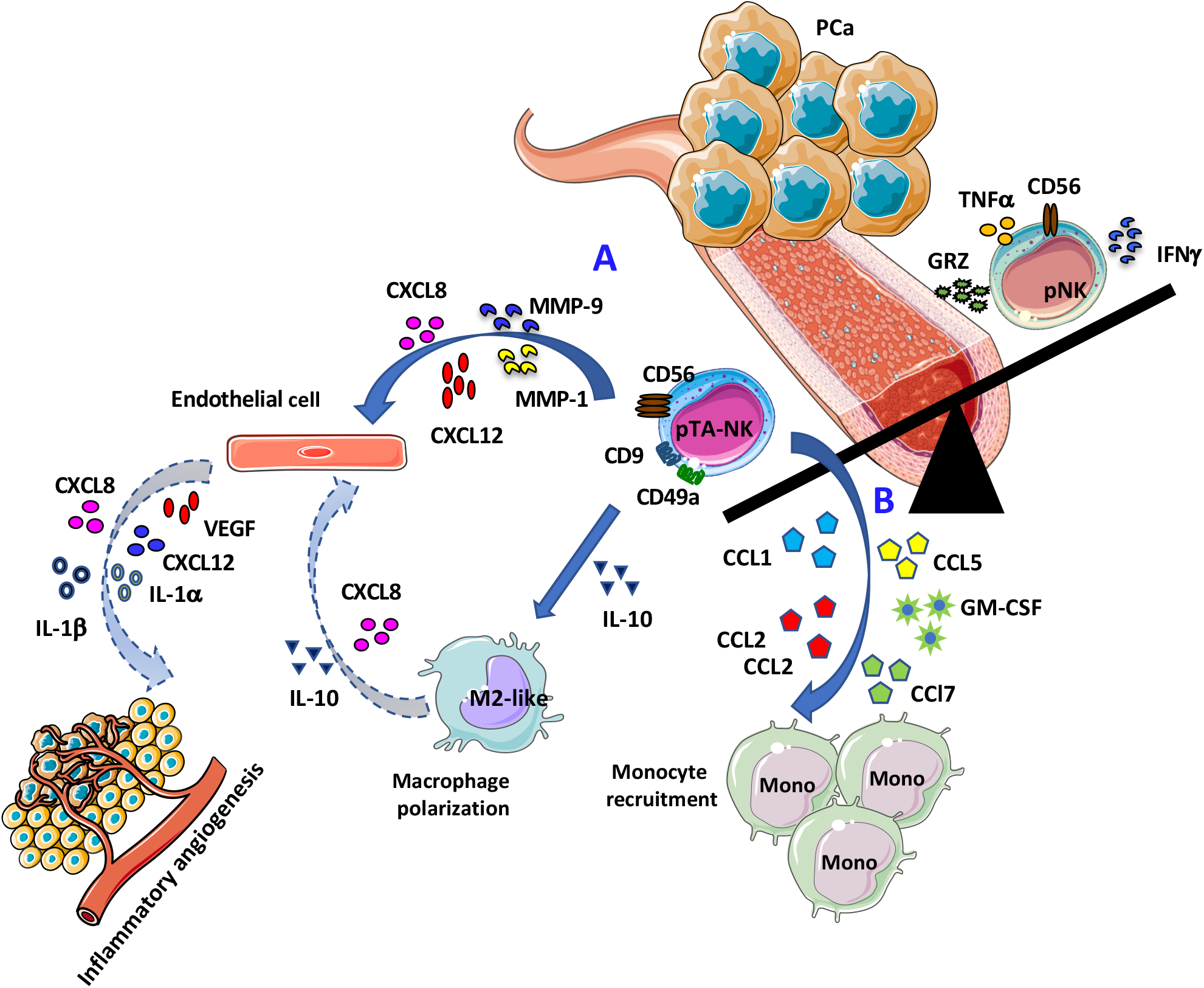

## INTRODUCTION

Prostate cancer (PCa), is the most frequent male cancer in the western world, and the second cause of cancer death in men [1]. Surgery and radiation therapy are still important treatment options, as well as chemotherapy and hormonal therapy for hormone sensitive cases. Recently immunotherapy came of age as a possible effective strategy for PCa therapy.

Evasion from immune system surveillance and induction of an inflammatory microenvironment, are among host-dependent biological features, widely accepted as cancer hallmarks, as defined by Hanahan and Weinberg [2] and which also plays a role in prostate cancer. Based on their extreme cell plasticity, inflammatory cells from innate and adaptive immunity have been reported to acquire tumour-promoting phenotypes and functions, in cancer patients [3–7]. Acquisition of a tolerogenic state, anergy and induction of inflammatory angiogenesis are some of these aberrant functions [3–6,8].

Natural killer (NK) cells are large granular lymphocytes endowed with an inherent capability to kill virally infected, and malignant cells and also to modulate the immune system through their production of numerous cytokines and chemokines. NK cells constitute approximately 5–15% of circulating lymphocytes in healthy adults and therefore represent one of the three major lymphocyte population. Although lymphocytic in origin NK cells are considered part of the innate immune system, not needing antigen presentation for recognition. They exert effector functions that include cytotoxic activity and cytokine production, during antiviral and antitumor responses [9]. As for several immune cells [3–6,8], NK cells have been described to acquire a tolerogenic behaviour and to be altered in their cytotoxic activities [3–6,10–16], they represent still an under investigated cell population and only few studies demonstrated pro-inflammatory angiogenic cancer cell growth promoting activities in cancers [3,10–12,14]. Major mechanisms associated with impaired NK cell function in cancer patients, are downregulation of lytic perforin/granzyme production accompanied with reduction of degranulation capabilities, together with reduction of NKG2D (the major NK cell activation receptor) expression [17–19]. In prostate cancer, the ligands for NKp30 and NKp46 are expressed on NK cells from patients with primary tumours but not on benign prostate hyperplasia [20]. However, studies on the prostate-infiltrating/associated NK phenotype and functions remain limited [21]. Isolation of tumour-infiltrating immune components is challenging, due to the small size of prostate biopsies, and the absence of stromal compartments. A study by D. Olive showed that inherent and tumour-driven immune tolerance in the prostate microenvironment impairs NK cell antitumor activity [16].

Two major subsets of NK are mostly present in the peripheral blood (pNK): the cytotoxic CD56^dim^CD16^+^ NK cell subset, (90–95% of pNK) and the low cytotoxic, highly cytokine producing NK cell subset, CD56^bright^CD16^-/low^ [9].

Our research group has identified a new pro-angiogenic NK cell subset in non-small-cell lung carcinoma (NSCLC) described as CD56^bright^CD16^-^VEGF^high^PlGF^high^CXCL-8^+^IFNγ^low^ NK cells [5,11,12], giving a role to NK cells in the inflammatory pro-angiogenic switch in solid tumours. These type of NK are similar to a peculiar NK subset that has been found within the developing decidua, the decidual NK cell (dNK) which exhibit a CD56^superbright^CD16^-^CD9^+^CD49a^+^ phenotype and are closely linked with vascularisation of the decidua and embryo implantation, in both humans and mice [22,23]. dNK cells produce VEGF, PlGF, and CXCL8, are poorly cytotoxic and are associated with induction of CD4^+^ T regulatory (Treg) cells [22,23]. We characterised also NK cells in the peripheral blood (tumour associated NK cells, pTA-NKs) and tissue infiltrate (tumour-infiltrating NK cells, TI-NKs) in colorectal cancer patients, these cell subsets also display pro-angiogenic feature as those in NSCLC patients [12]. NK cells in the peripheral blood, in particular the CD56^bright^CD16^low/-^, reflect the TI-NKs, and are present both in male and female cancer patients [11,12,24,25].

We identified TGFβ, a major immunosuppressive cytokine in the TME [26,27], as an inducer of the inflammatory pro-angiogenic switch of cytolytic NK cells both at tissue and peripheral levels [12]. We found that Stat3/Stat5 activation regulates the polarization in CRC NK cells and that Stat5 chemical inhibition, interferes with this process [10,25]. Several other groups have reported features or polarize pNK in tumors, however PCa still needs more investigations.

Here, we show for the first time that NK cells isolated from peripheral blood of patients with PCa (PCa pTA-NKs), acquire a pro-inflammatory angiogenesis phenotype, characterized by increased expression of the surface antigens CD56, CD9 and CD49a. Analysis on secreted products by FACS-sorted NK from PCa blood samples, allowed the identification of three main pNK signatures, characterized by up-regulation of cytokines and chemokines with pro-inflammatory angiogenesis/pro-metastatic features (CXCL8/IL-8, MMP-1, MMP-9, uPAR, TIMP-1), promonocyte recruiting features (CCL1, CCL2/CCL5) and properties involved in M2-like macrophage polarization (IL-10). Secreted products of FACS-sorted pNK cells from peripheral blood of PCa patients were able to polarize THP-1 macrophages towards *CD206/Arginase1/CXCL8/IL10*-expressing M2-like/TAMs. Differently than in our previous studies, we could not identify large numbers of CD56^superbright^CD16^-^ NK cells in the prostate cancer tissue (data not shown). However increasing evidence suggests that polarized/polarizing NK cells are not restricted to those infiltrating the tumour, but are found to be present in peripheral blood of patients with several types of cancer [4,10–12,24,25,28], where they could represent a valuable independent marker.

Our results contribute in placing circulating pNK cells as important innate immune pro-inflammatory effector cells that could represent a biomarker in prostate cancer patients.

## MATERIALS AND METHODS

### Samples selection and patient characteristics

Peripheral blood (PB) samples (15-20 mL) were obtained from patients with prostate adenocarcinoma (ADK, n=25). Controls (HC, n=20) included peripheral blood of healthy, tumour-free, individuals. Patients with diabetes, human immunodeficiency virus (HIV)/hepatitis C virus (HCV)/hepatitis B virus (HBV) infection, chronic inflammatory conditions, treated with chemotherapy or radiotherapy, iatrogenically immunosuppressed or subjected to myeloablative therapies, were excluded to the study. The study was approved by the institutional review board ethics committees (Protocol N°0024138 04/07/2011 and Protocol N°10 2 10/2011, within the study PROSTATEST) and, according to the Helsinki Declaration of 1975 as revised in 2013. All patients enrolled in the study signed the informed consent, in accordance to the Helsinki Declaration of 1975 as revised in 2013. Demographic features of the cohort of PCa patients and controls are showed in Supplementary Table 1.

### Cell culture and maintenance

The human prostate cancer (PCa) cell lines PC-3, DU-145, LNCaP (all purchased by ATCC) were maintained in in RPMI 1640 medium, supplemented with 10% Fetal Bovine Serum (FBS), (Euroclone), 2 mM l-glutamine (Euroclone), 100 U/mL penicillin and 100 μg/mL streptomycin (Euroclone), at 37°C, 5% CO_2_. Cells were routinely screened for eventual mycoplasma contaminations. Conditioned media (CM) were collected following 72 hours of starvation in FBS free RMPI 1640 (Life Technologies), supplemented with 1% Glutamine (Euroclone) and 1% P/S (Euroclone), at 37°C, 5% CO_2_. CMs were used for NK cell polarization as detailed below.

Human umbilical vein endothelial cells (HUVEC, Lonza) were maintained in endothelial cell basal medium (EBM™, Lonza) supplemented with endothelial cell growth medium (EGM™ SingleQuots™, Lonza), 10% of FBS, 2 mM l-glutamine (Euroclone), 100 U/mL penicillin and 100 μg/mL streptomycin (Euroclone). HUVEs were used between the 3–5 passages.

The human monocytic THP-1 cell line (ATCC) was cultured in RPMI 1640 medium, supplemented with 10% FBS, 2 mM l-glutamine (Euroclone), 100 U/mL penicillin and 100 μg/mL streptomycin (Euroclone), at 37°C, 5% CO_2_. Differentiation of adherent THP-1 macrophages was obtained following 48 hours of treatments with phorbol-merystate-acetate (5 ng/mL, PMA, Sigma Aldrich), as in [29].

### NK cell isolation by FACS-sorting

pNK cells were isolated from peripheral blood mononuclear cells (PBMCs) of PCa -ADK and HC subjects. Whole blood samples (20 mL) were diluted with PBS 1:1 (v/v), then subjected to a density gradient stratification with Ficoll Histopaque-1077 (Sigma-Aldrich), at 500g for 20 minutes. The white ring, composed of total mononuclear cells (MNCs), was collected, washed twice in PBS, then used for subsequent experiments or for pNK isolation. Total MNCs were subject to cell sorting, using a BD FACS-AriaII instrument. For details of antibodies used, see Supplementary table 3. Following 30 minutes of staining with anti-human PerCP conjugated CD3 and anti-human APC-conjugated CD56, NK cells was sorted as lymphocyte (SSC/FSC) gated CD3^-^CD56^+^ cells. Post sorting purity was immediately checked. Purified NK cells (2*10^5^ cells/mL) were used, following 24 hours of culture in serum-free RPMI, for molecular analysis (qPCR) and to collect conditioned media for functional and secretome studies. After 24 hours, supernatants were collected, centrifuged to remove residual dead cells and debris and concentrated using Concentricon (Millipore) with a 3kDa membrane pore cut-off to obtain concentrated supernatants.

### Cell treatment with secreted products

For NK cell polarization, total PBMCs (1*10^6^ cells/mL) were polarized with 30% of PC-3 or DU-145 or LnCaP CMs (v/v) in RMPI 1640 (Euroclone), supplemented with 10% FBS (Euroclone), 2 mM l-glutamine (Euroclone), 100 U/mL penicillin and 100 μg/mL streptomycin (Euroclone), 100 U/mL IL-2 (R&D), at 37°C, 5% CO_2_, for 72 hours. Cells were pulsed with fresh CMs and complete RPMI (30%, v/v) at day 0 and at 48 hours, during the polarization schedule.

Conditioned media from FACS-sorted NK cells were used to detect the production of pro-pro-inflammatory factors by endothelial cells. 2×10^5^ HUVE cells were seeded into six well plates ant exposed for 24 hours to CM (50 μg/mL of total protein) of PCa pTA-NKs or NK cell from HC. HUVECs were then harvested and used for real-Time PCR analysis.

THP-1 differentiated macrophages were obtained from THP-1 monocytes treated for 48 hours with 5 ng/mL of PMA. Following attachment, THP-1 differentiated macrophages were pulsed with CMs (50 μg of total protein) from FACS-sorted NK cells (either from PCa patients of controls) for 72 hours. THP-1 macrophages received CM at day 0 and 48 hours of stimulation. Expression of M1-like or M2-like/TAM markers, following polarization, was detected by Real-Time PCR.

### Phenotype characterization of CM-polarized and PCa patient derived NK cells

The polarization state of either pNK cells exposed to PCa cell line (PC-3, DU-145, LNCaP) conditioned media or pNK cells from PCa patients, was assessed by flow cytometry for surface antigen expression. Briefly, 2.5×10^5^ of total PBMCs per FACS tube were stained for 30 minutes at 4°C with anti-human monoclonal antibodies (mAbs) as follows: PerCP conjugated anti-CD3, APC conjugated anti-CD56, FITC conjugated anti-CD16, PE conjugated anti-CD9, PE conjugated anti-CD49a, PE conjugated anti-NKG2D (all purchased by Miltenyi Biotec). Following Forward/Side Scatter setting, NK cells were identified as CD3^-^ and CD56^+^ cells (total NK cells). CD16 and NKG2D expression was evaluated on CD3^-^CD56^+^ (total NK) gated cells. Finally, CD56 brightness and the expression of the dNK markers CD9, CD49a and CXCR4, were evaluated on total CD3^-^CD56^+^NK cells. For details on antibodies used, see Supplementary table 3.

### Intracellular staining for cytokine detection of CM-polarized pNK cells

For intracellular cytokine detection, 2×10^6^ PBMCs from PCa-ADK patients or HC were cultured, overnight, in RPMI 1640 (EuroClone) supplemented with 10% FBS (Life Technologies,) 1% (v/v) L-Glutammine (Sigma), 100 U/mL penicillin, 100 μg/mL streptomycin (Euroclone) and IL-2 (100 U/ml; R&D Systems) at 37°C and 5% CO_2_. For intracellular staining, the third day of polarization, cells were stimulated for 6 h with PMA (10 ng/mL) and ionomycin (500 ng/mL) (both from Sigma), in the presence of GolgiStop plus GolgiPlug (both from BD Biosciences).

Cells were collected and stained for NK cell surface markers, as previously described [ref], washed with PBS and treated with Cytofix/Cytoperm fixation and permeabilization kit (BD) for 10 minutes at 4°C. Cells were then washed in PBS and stained with PE-conjugated anti human VEGF, CXCL8, CXCL12 (R&D System), IFNγ, TNFα, GranzymeB (Myltenyi Biotec) for 30 minutes. For indirect staining, cells were incubated for 1 hour at 4°C with primary unlabelled antibodies antihuman Angiopoietin 1, anti-human Angiogenin, (all purchased from Abcam), washed and then stained with secondary PE-conjugated labelled antibody anti-mouse IgG, for 30 minutes, at 4°C. Cytokines production was detected by flow cytometry, using the BD FACS CantoII analyzer. Isotype control and the secondary antibody alone were used as staining controls. For details on antibodies used, see Supplementary table 3.

### Secretome analysis of PCa pTA-NKs

The secretome of conditioned media (50 μg of total protein) of FACS-sorted pNK was assessed, using the Human Angiogenesis Array C1000 (RayBiotech, Inc., Norcross GA) to detect cytokines and chemokines release, as in [25]. A pool of three ADK or HC were used. Chemiluminescent signals (revealed as black spots) were captured by membrane exposure to Amersham Hyperfilm. Arrays were computer scanned using the Amersham Imager 680 Analyzer and optical density was determined using the ImageJ software.

### Network formation assay on endothelial cells

HUVEC cells (1,5 x10^5^ cells/well) were seeded in a 96 well plate, previously coated with 50 μL of 10 mg/mL polymerized Matrigel (BD). After exposure to secreted products (50 μg/mL total protein), in serum-free EBM medium, HUVECs were then incubated at 37°C, 5% CO_2_ for 24 hours. The formation of capillary-like structures was detected by microphotographs, using an inverted microscope (Zeiss). The number of master segment and master segment length, as indicators of tube formation efficiency, were determined, using ImageJ software and the Angiogenesis Analyzer tool.

### Detection of THP-1 monocyte recruitment by PCa pTA-NKs

Migration assay was performed using modified Boyden chambers. 5 x 10^4^ THP-1 monocytes were resuspended in 500 μL of serum-free RPMI and loaded into the upper compartment of the Boyden chamber. The lower chambers were filled with 250 μL of serum-free RMPI medium, supplemented with 50 μg of total protein of secreted products from ADK or HC pNK cells. 3 μm pore-size polycarbonate filters (Whatman, GE Healthcare Europe GmbH, Milan, Italy) previously pre-coated with 2 μg/mL of fibronectin, were used as interface between the two chambers. The Boyden chambers were incubated for 6 hours at 37°C. Filters were recovered, cells on the upper surface mechanically removed with a cotton swab. Cells migrated toward the lower filter surface, were fixed with ethanol at serial percentage (70%, 100%), finally rehydrated in water. Filters were stained with 10 μg/mL DAPI (Vectashield, Vector Laboratories, Orton Southgate, Peterborough, United Kingdom) and incubated at room temperature, protected from light for 10 minutes. Cells in the filters were counted in a double-blind manner in five consecutive fields/filter, with a fluorescent microscope (Nikon Eclipse).

### Quantitative Real-time PCR (qRT-PCR)

Total RNA was extracted from FACS-sorted pNK cell, CM-exposed HUVECs or THP-1 macrophages, using the small RNA miRNeasy Mini Kit (Thermo Fisher) and quantified by Nanodrop Spectrophotometer. Following genomic DNA removal, by DNase I Amplification Grade (Thermo Fisher) treatment, reverse transcription was performed on 500 ng of total RNA using SuperScript VILO cDNA synthesis kit (Thermo Fisher). Real-time PCR was performed using SYBR Green Master Mix (Thermo Fisher) on QuantStudio 6 Flex Real-Time PCR System Software (Applied Biosystems, Thermo Fisher Scientific, USA). All reactions were performed in triplicate. The relative gene expression for pro-angiogenic factors (in HUVECs) or M1/M2-like markers (in THP-1 macrophages) was expressed relative to healthy controls, normalized to GAPDH ([CT (gene of interest) −CT(GAPDH)] = ΔCT). HUVECs or THP-1 macrophages in their respective basal medium alone, were used as baseline controls. Primer sequences are provided in Supplementary Table 2.

### Statistical analysis

Statistical differences between two datasets were determined using two tailed t-test. For multiple datasets, analysis of variance (ANOVA) followed by Tukey’s post-hoc test was used. P values (p) ≤ 0.05 will be considered statistically significant. Data were analysed using the GraphPad Prism8 (San Diego, CA). Flow cytometry data were analysed using the BD FACS-Diva and FlowLogic software.

## RESULTS

### pTA-NKs from PCa patients exhibit a decidual-like, pro-inflammatory phenotype

We investigated whether pNK from PCa patients are characterized by a pro-inflammatory angiogenesis phenotype. Flow cytometry analysis of CD56 and CD16 surface antigen expression revealed that the CD56^+^CD16^+^ NK cells are the predominant subset in the peripheral blood in PCa-ADK and HC samples (Supplementary Figure 1A). We found increased frequency of CD56^bright^ NK cells in the peripheral blood of patients with PCa ADK (***p ≤ 0.001), (Figure 1A). Peripheral blood NK cells from PCa-ADK samples express also higher levels of the decidual-like markers CD9 (Figure 1B), CD49a (Figure 1C) as compared with those isolated from healthy controls (*p ≤ 0.05; **p ≤ 0.01; ***p ≤ 0.001). We also found increased expression of CXCR4 on NK cells from PCa-ADK samples (Figure 1D). The NKG2D expression on PCa TA-NKs was reduced (*p ≤ 0.05), as compared to those from HC (Supplementary Figure 1B). Real-time PCR results showed that TA-NKs cells, FACS-sorted from PCa-ADK, have increased expression of RNA for the pro-inflammatory factors *CXCL8* (**p ≤ 0.01), *CXCL12* and *PAI* and confirmed the increased RNA expression of *CXCR4*, as well as *VEGF* (****p ≤ 0.0001), as compared to NK isolated form the peripheral blood of healthy controls (Figure 2E).

**Figure 1.**
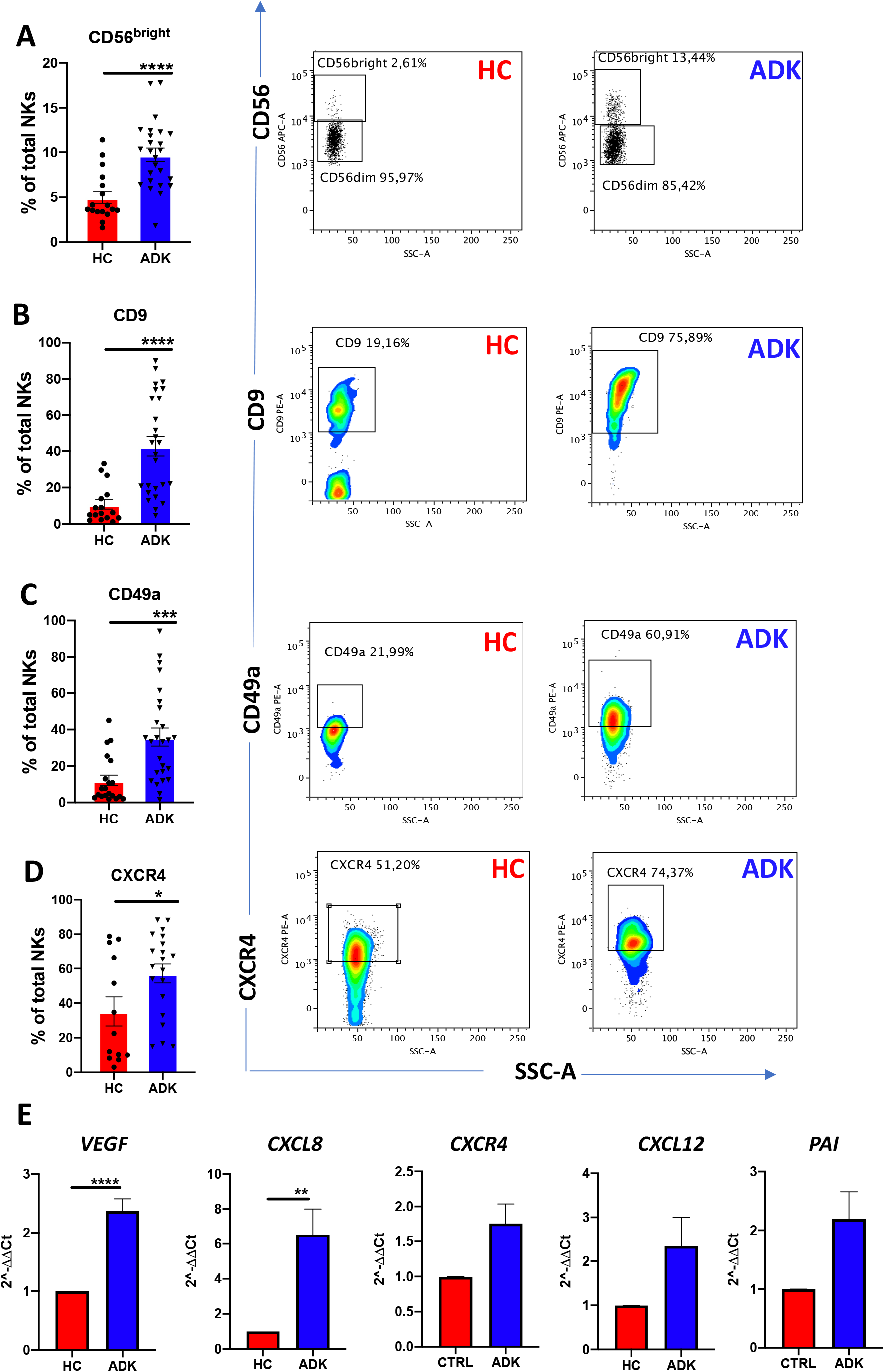
pNK cell polarization in peripheral blood of PCa patients. PCa TA-NKs have increased frequency of CD56^bright^ NKs as compared with those from HC (A). Peripheral blood NK (pTA-NKs) from PCa patients significantly express higher levels of the dNK cell markers CD9 (B) CD49a (C) and CXCR4 (D), as compared with those from HC. Representative dot plots show the specific antigen expression (as percent on total pNK cells). pNK cells FACS-sorted from patients with ADK-PCa have increased expression of the pro-inflammatory factors *VEGF, CXCL8, CXCR4, CXCL12, PAI* (E). qPCR have been performed using pNK cell from 6 PCa patients and 6 PCa controls, in triplicate. Data are showed as mean ± SEM, t-student test, *p<0.05, *p<0.01, ***p<0.001, ****p<0.0001. HC: healthy controls; ADK: prostate cancer adenocarcinoma.

**Figure 2.**
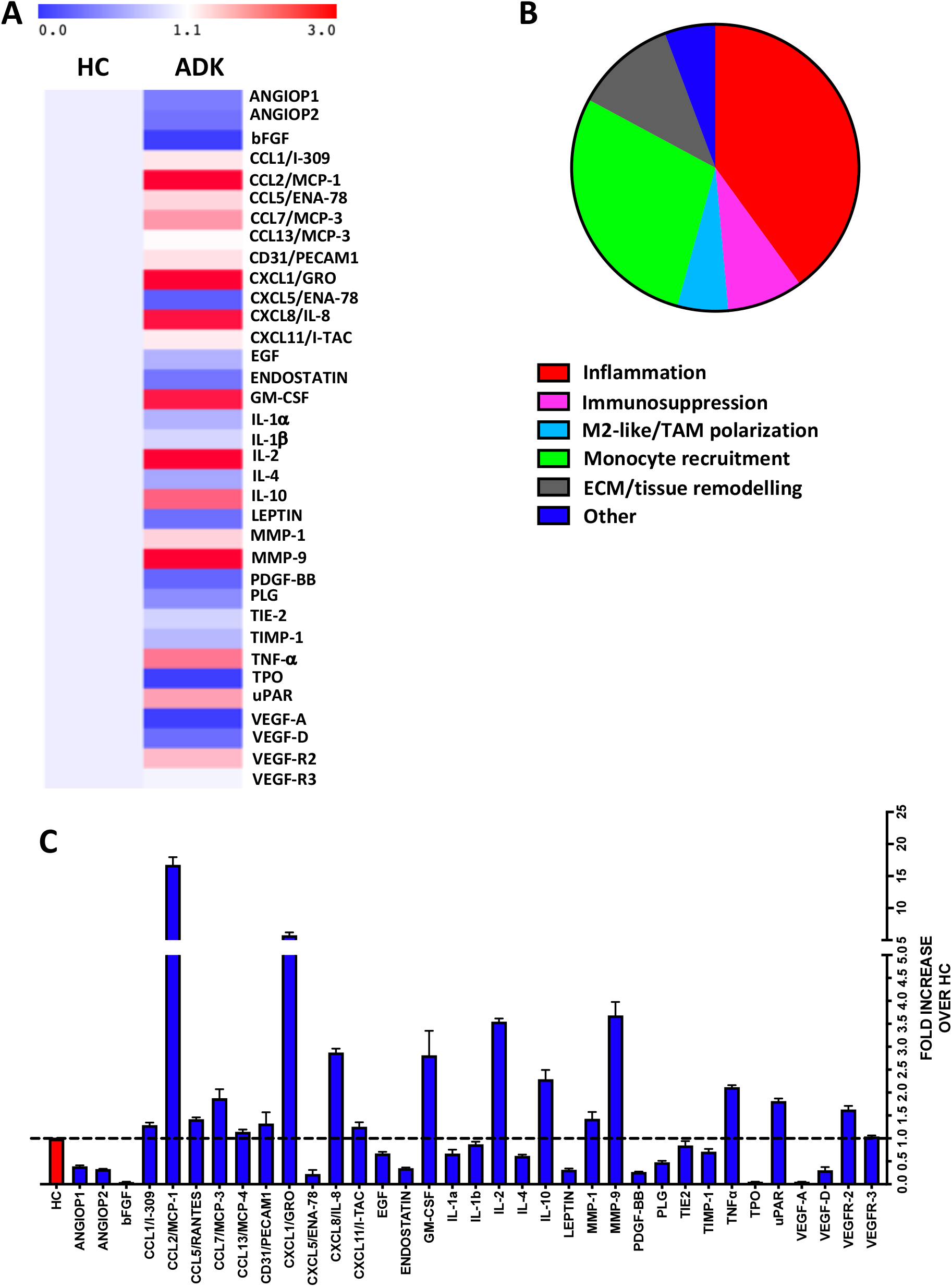
Secretome profiling of PCa pTA-NKs. Secretome profiling, using an antibody membrane array showed that conditioned media from PCa patients are enriched in several factors directly and indirectly associated with induction of angiogenesis, immunosuppression, M2-like polarization, macrophage recruitment, ECM/tissue remodelling. A) Representative heatmap; B) part of whole diagram showing the frequency for the signatures observed; C) bar histogram showing the fold change for every modulated factor PCa/HC. CMs were pooled from NK cells sorted from 3 different PCa patients. Arrays were performed in duplicate. HC: healthy controls; ADK: prostate adenocarcinoma.

### pTA-NKs from PCa patients can release pro-inflammatory cytokines and chemokines involved in monocyte recruitment and polarization

To investigate whether the acquisition of the pro-inflammatory phenotype in PCa pTA-NKs would correlate with their capability to release soluble factors involved in direct and indirect induction of inflammatory-angiogenesis, we investigate the contents of CM from PCa TA-NKs. We characterized the production of secreted proteins from PCa pTA-NKs using a commercially available angiogenesis-membrane array kit. The overall secretome analysis (Figure 2A-C, Supplementary Figure 2) revealed three major signatures characterizing PCa-ADK pTA-NKs: tissue inflammation/remodelling signature (CXCL8, MMP-1, MMP-9, uPAR, TIMP-1) a monocyte recruiting signature (CXCL1, CCL2, as the most up-regulated) and M2-like macrophage polarization (Figure 2A-C).

### pTA-NKs from PCa patients functionally support inflammatory angiogenesis *in vitro*

We further investigated whether PCa-ADK pTA-NKs, expressing pro-inflammatory cytokines (also involved in angiogenesis), chemokines and chemokine receptors, were also effectively able to induce network formation in HUVECs *in vitro*. We found that conditioned media (CM) of pNK cells isolated PCa-ADK samples have higher contents of the pro-inflammatory/tissue-remodelling factors CXCL8/IL-8 (****p≤0.0001), MMP-1 (*p≤0.05), MMP-9 (****p≤0.0001), uPAR (****p≤0.0001) (Figure 3 A). To detect whether secreted products of inflammatory NK cells isolated PCa-ADK samples were effectively able to induce network formation in HUVECs, we treated HUVE cells with these secreted products. We found that secreted products of pNK cells isolated from PCa-ADK are able to induce the formation of capillary-like structures by HUVE cells, on a matrigel layer (*p ≤ 0.05; **p ≤ 0.01), as a consequence of their pro-inflammatory secretome (Figure 3B). Real-time PCR results showed that HUVE cells, exposed for 24 hours to CM from PCa-ADK pTA-NKs have a pro-inflammatory phenotype with increased expression of *VEGF* (*p ≤ 0.05), *VEGF-R2, CXCL8* and of factors involved in vascular inflammation and immune cells mobilization, such as *CXCR4/CXCL12* axis, *ICAM-1, VCAM-1*, together with induction of *IL-1*α (***p ≤ 0.001) (Figure 3C).

**Figure 3.**
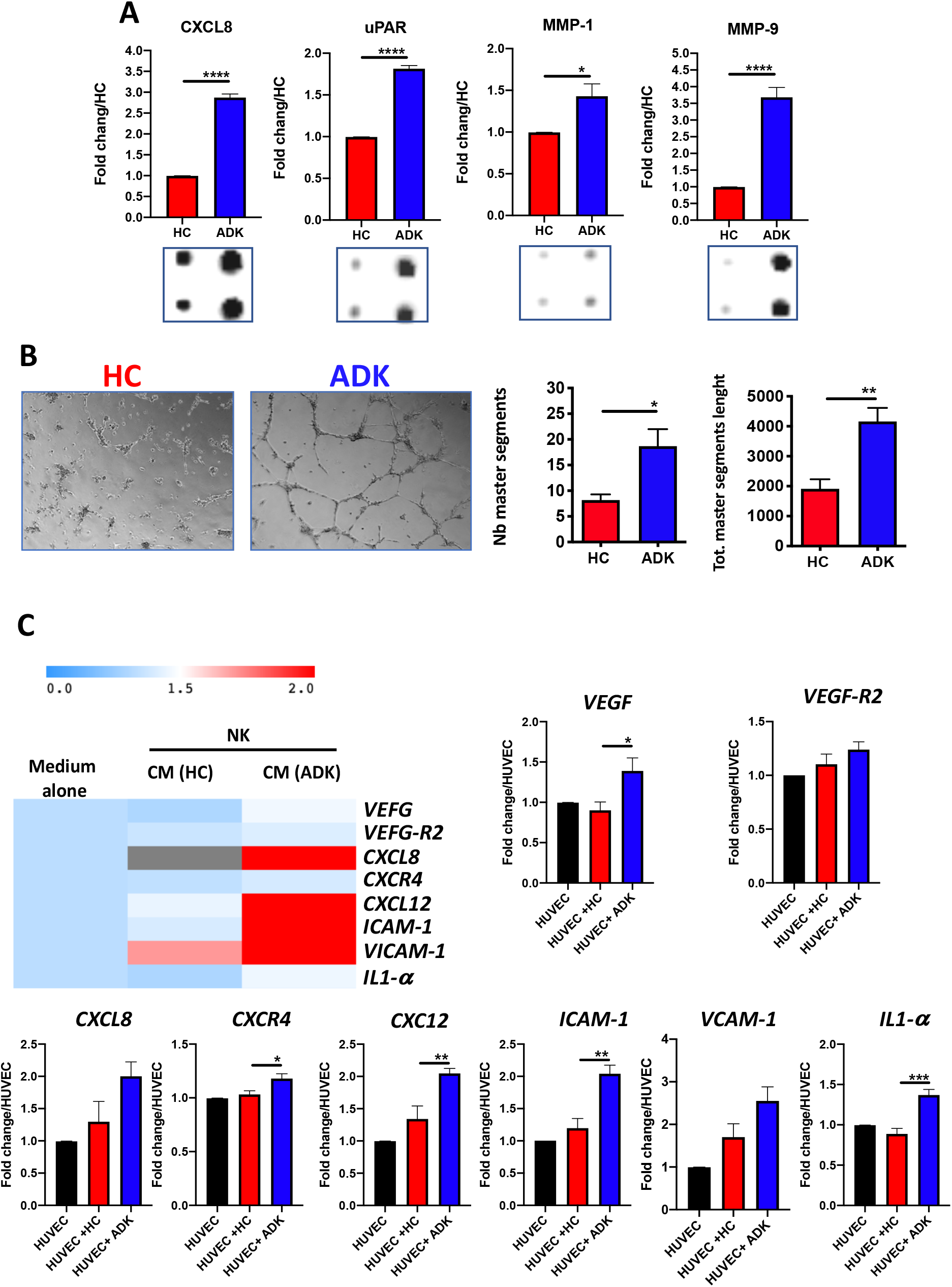
Pro-inflammatory activities of pTA-NKs from PCa patients on endothelial cells. Secreted products (CM) form FACS-sorted PCa pTA-NKs are enriched in pro-inflammatory and tissue-remodelling factors, such CXCL8, uPAR, MMP-1, MMP-9 (A) and functionally support the formation of capillary like structures in human umbilical-vein endothelial cells (HUVEC) on matrigel (B). HUVE cells exposed to secreted products of PCa pTA-NKs express higher levels of pro-inflammatory factors like *VEGF, VEGF-R2, CXCL8, CXCR4, CXCL12, ICAM-1, VCAM-1, IL1-α*, as compared to those exposed to secreted products released by healthy control NK cells (C). Capillary like-structure formation and qPCR on HUVECs have been performed using CM of pNK cell from 6 PCa patients and 6 PCa controls, in triplicate. Data are showed as mean ± SEM, ANOVA, *p<0.05, **p<0.01, ***p<0.001, ****p<0.0001. The condition HUVEC (black bar) stands for HUVE cells alone, as baseline condition. HC: healthy controls; ADK: prostate adenocarcinoma.

### pTA-NKs from PCa patients can recruit monocytes and induce an M2-like/TAM features

Secretome analysis revealed that secreted products of pNK cells isolated PCa-ADK samples are enriched in soluble factors involved in macrophage recruitment and polarization (Figure 2, Supplementary Figure 2) with GM-CSF (*p ≤ 0.05), CXCL1/GRO (****p ≤ 0.0001), CXCL11/I-TAC (*p ≤ 0.05), CCL1/I-309 (**p ≤ 0.01), CCL2/MCP-1 (****p ≤ 0.0001), CCL5/RANTES (****p ≤ 0.0001), CCL7/MCP-3 (**p ≤ 0.01), CCL13/MCP-4 (*p ≤ 0.05) and IL-10 (***p ≤ 0.001) (Figure 5A). Based on these results, we functionally investigated PCa-ADK NK cells ability to recruit macrophages, via soluble factors. We observed that CM of pNK cells purified from the peripheral blood of PCa-ADK patients, significantly (***p ≤ 0.001) promote the recruitment of THP-1 monocytes, as compared to CM of NK cells isolated from healthy controls (Figure 4B). We also observed that THP-1 differentiated macrophages, following 72 hours of exposure to ADK pTA-NK CMs, displayed a decreased expression of the M1-like cytokine IL-12 (*p ≤ 0.05) and an increased expression of the M2-like/TAM factors; CD206/Mannose receptor, Arg1 (*p ≤ 0.05), CXCL8 (***p ≤ 0.001) and IL-10 (**p ≤ 0.01) (Figure 4C).

**Figure 4.**
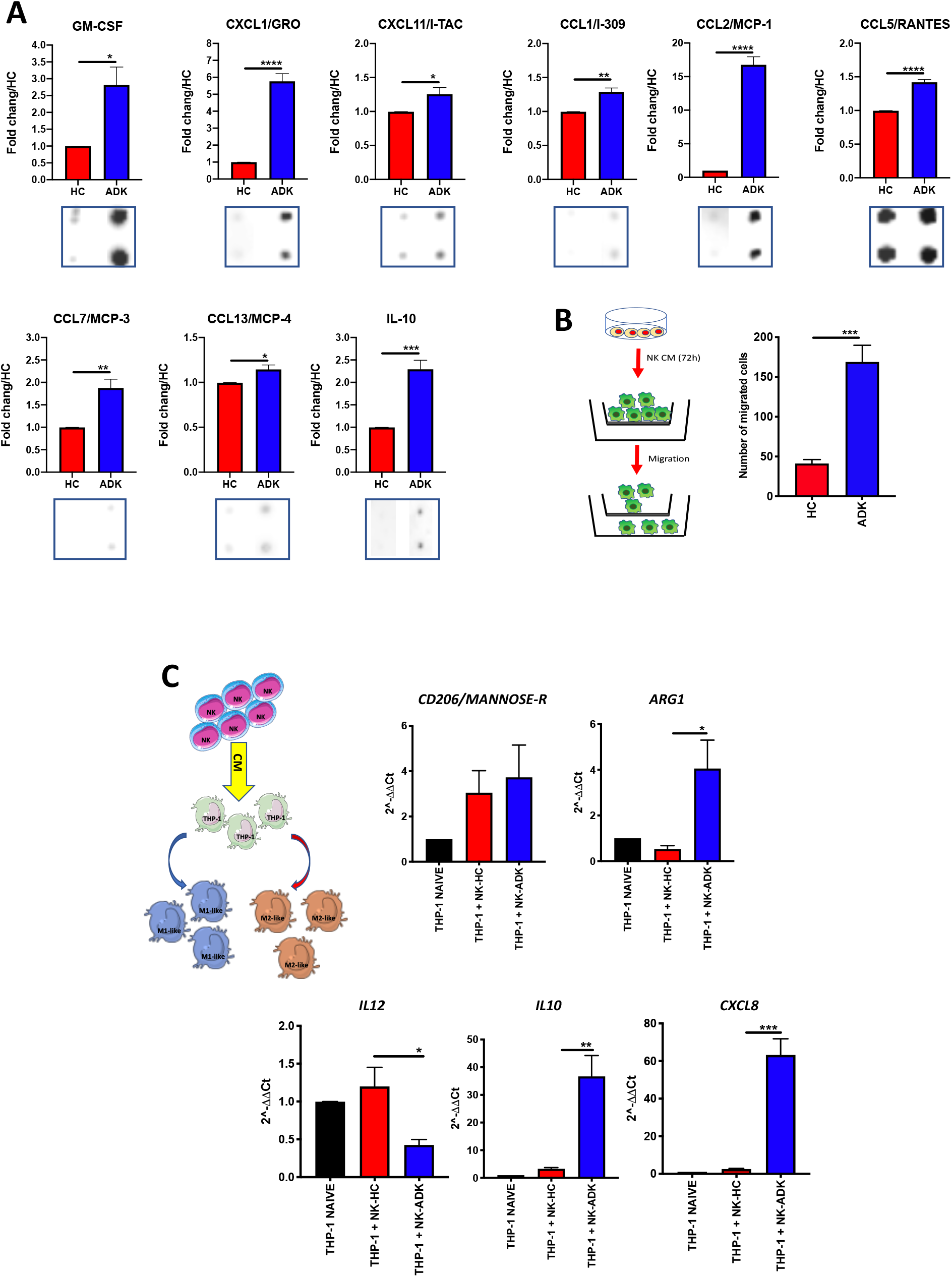
Effects of PCa pTA-NKs on monocyte recruitment and polarization. Conditioned media from PCa pTA-NKs are enriched with factors involved in macrophage recruitment (GM-CSF, CXCL1/GRO, CXCL11/I-TAC, CCL1/I-309, CCL2/MCP-1, CCL5/RANTES, CCL7/MCP-3, CCL13/MCP-4, and polarization (IL-10) (A) and can recruit THP-1 monocytes as compared with those from heathy controls (B), as revealed by the migration assay (Boyden Chambers) (A). Exposure of THP-1 activated macrophages to conditioned media of pNK from PCa patients results in THP-1 cells ability to express higher levels of M2-like/TAM markers (*IL-10, ARG1, CXCL8*) while reducing the expression of *IL-12* (M1-like marker) (C). CMs were pooled from pNK cells FACS sorted from 3 different PCa patients. Arrays were performed in duplicates. Data are showed as mean ± SEM, ANOVA, **p<0.01, ***p<0.001, ****p<0.0001. HC: healthy controls; ADK: prostate adenocarcinoma; naive indicates THP-1 cells in control medium.

**Figure 5.**
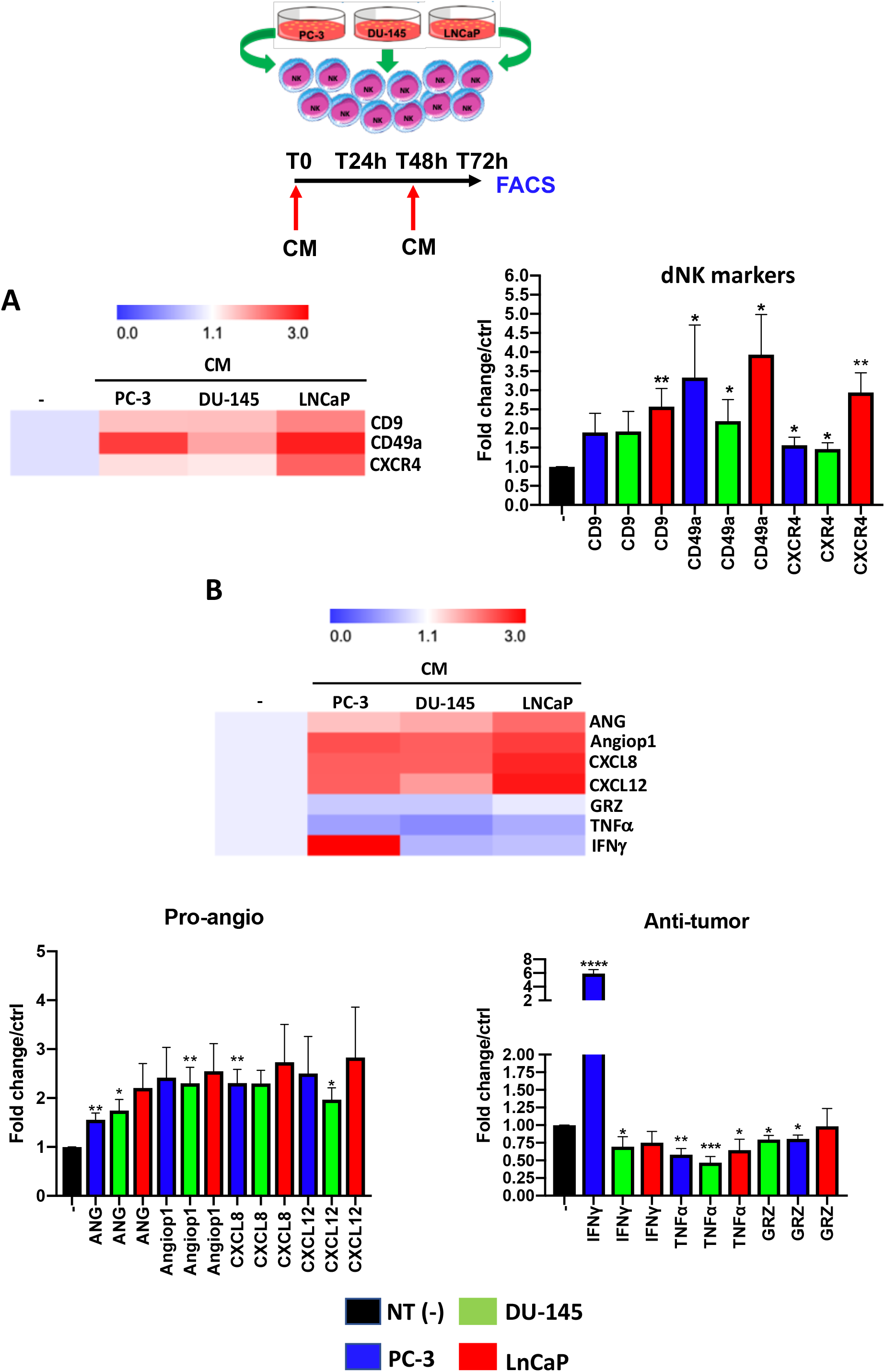
Effects of prostate cancer cell line secreted products (CM) on pNK cell polarization. NK cells from healthy donors, following 72 hours of exposure to conditioned media (CM) from PC-3, DU-145 and LNCaP PCa cell lines, exhibit a pro-inflammatory angiogenic decidual-like phenotype, as revealed by the increased levels of the dNK-like markers CD9, CD49a, CXCR4 (A), enhanced production of pro-inflammatory factors (angiogenin, ANG; angiopoietin-1, Angiop1; CXCL8) and reduced production of cytolytic factors (granzyme B, GRZ-B; TNFα; IFNγ), as revealed by flow cytometry analysis (B). Experiments were performed using peripheral blood samples of 5-to-9 independent healthy donors. Data are shown as mean ± SEM, ANOVA, *p<0.05, **p<0.01, **p<0.001. CM: conditioned media/secreted products from 72 hours of SFM PCa cell lines.

### Prostate cancer cell lines secreted products polarize pNK cells towards pro-inflammatory angiogenic NK cells

We used an *in vitro* model mimicking the interaction of the secretome of PCa with normal PBMC of healthy donors. Mononuclear cells from peripheral blood were exposed to soluble factors (conditioned media, CM) collected from three different PCa cell lines (PC-3, DU145, LNCaP) and assessed for their expression of decidual and pro-inflammatory, angiogenic markers and polarization state. We found that pNK cells from healthy controls, following 72 hours of exposure to the CM of three different PCa cell lines (PC-3, DU-145, LNCaP) showed increase expression of the CD9, CD49a decidual markers and of CXCR4 (*p ≤ 0.05, **p ≤ 0.01) chemokine receptor (Figure 5A). We also found that 72 hours of stimulation with CM from the three PCa cell lines resulted in pNK ability to produce enhanced chemokines CXCL8 (**p ≤ 0.01) and CXCL12 (*p ≤ 0.05) as detected by flow cytometry (Figure 5B).

## DISCUSSION

Although immunotherapy has emerged as the “next generation” of cancer treatment [30], it has not yet been shown to be always successful in the treatment of patients with PCa, for whom the preferential therapeutic options still remain radiotherapy, chemotherapy, androgen deprivation therapy [31–34]. This clearly suggest that, to address more efficient (immune)therapeutic approaches against PCa, a better understanding of how the PCa is able to subvert the host immune system, still remains a major issue and a clinical unmet need. Preclinical and clinical evidences suggest that chronic inflammation plays a crucial role in multiple stages of prostate cancer development [35–37].

The polarization of the immune inflammatory cells in peripheral blood is directed by specific chemokines and cytokines that can shape their state and make them acquire altered phenotype/functions, depending on tumour scenario [3,5,6].

NK cells have been found to be compromised in several cancers [3,6,9–13,15,18,19,24,25,38]. NK cell contribution to tumour progression goes beyond the mechanism of tumour escape and immunosuppression [3,10–12,24,25]. We demonstrated that, NK cells in NSCLC cancer [12], colorectal cancer [3] and in malignant pleural effusions [24], show a pro-angiogenic, pro-inflammatory phenotype and functions, identified as CD56^bright^CD16^-^VEGF^high^CXCL8^+^IFNγ^low^ and share several features/behaviours with the highly pro-angiogenic dNK cells. This was confirmed by other groups in breast and colon cancers [39].

Very little is known about NK alterations in prostate cancer, pNK scenario in PCa is less investigated. Here we studied pNK cells isolated from the peripheral blood of patients with PCa, in the framework of an approved clinical protocol. We found that peripheral blood NK cells from PCa patients acquire the CD56^bright^CD9^+^CD49a^+^CXCR4^+^ phenotype, indicating an inflammatory polarization, and we hypothesize that circulating pNK cells could serve as a biomarker for PCa patients. We show here that circulating pNK cells from PCa patients were able to express larger amount of the pro-inflammatory factors CXCL8, CXCL12, PAI, as compared to those from controls. We asked whether soluble-related factors, released by PCa pTA-NKs, might support inflammatory-like behaviour, acting on cellular components of the innate immune system, such as monocyte/macrophages. NK cells can interact with most of the innate and adaptive cellular components of the immune system [3,6,10,40,41]. Monocytes are the second most represented phagocytes in circulation and in established progressing tumours, were they display an M2-like/TAM phenotype [42–44] *In vitro* induced M2-like macrophages have been shown to decrease the susceptibility of tumour cells to NK cell cytotoxicity, with increased programmed death receptor ligand 1 (PD-L1) and decreased NK group 2D (NKG2D) ligands in castration-resistant prostate cancer (CRPC) cells [45].

Here we analysed the PCa pTA-NKs production of pro-inflammatory factors, using commercially available protein membrane arrays. We were able to distinguish three different secretome signatures that are not only restricted to endothelial cell regulation in angiogenesis. We found elevated release pro-inflammatory/tissue-remodelling cytokines and factors, of CXCL8/IL-8, MMP-1, MMP-9, uPAR, which can be responsible for the pTA-NK soluble-factor mediated induction of capillary-like structures on matrigel by HUVE cells. These results support the hypothesis that pNK cells from PCa patients can promote inflammatory angiogenesis, thus contributing to prostate cancer progression. A number of studies have linked CXCL8/IL-8 higher serum levels or expression in TME with aggressive prostate cancer, high Gleason score and with AR loss in metastatic disease [46–49]. Other studies reported that MMP-1, MMP-9, uPAR play important roles in tissue remodelling with prognostic implication PCa [50–52]. In previously published results, we reported that MMP-9 is upregulated in peripheral blood NK cells of colon cancer patients and the TIMP1/MMP9 axis, as well as uPAR, are altered as compared to normal circulating NK cells [25].

The crosstalk between NK cells and M1 macrophages plays a crucial role in the protection against infections and tumour development [53–55]. Tumour-derived monocytes induce rapid, transient activation, but subsequent, dysfunction and early apoptosis in human NK cells [55]. Conditioned media (CM) of M2 type macrophages decreases the susceptibility of tumor cells to NK cell cytotoxicity, as a result of increased programmed death receptor ligand 1 (PD-L1) and decreased NK group 2D (NKG2D) ligands in prostate cancer cells, through the IL-6 and STAT3 pathway [45].

While macrophage-NK cell crosstalk has been investigated in different cancers [53,54,56–58], PCa TA-NKs are able to produce several cytokines and chemokines involved in monocyte recruiting and macrophage polarization. We assessed the ability of PCa pTA-NKs to recruit monocytes in an *in vitro assay*. We found that PCa pTA-NKs have increased ability to stimulate migration of THP-1 monocytes as compared to pNK cells from healthy controls. We therefore tested whether the PCa products may impact on macrophage polarization state. We found that THP-1-differentiated macrophage, exposed for 72 hours to conditioned media from PCa pTA-NK cells, produce IL10 and Arginase (M2-Like factors) and decreased IL-12 (M1-like cytokine), at transcript level. We also observed increased expression of CXCL8 in THP-1 cells exposed to the same secreted products. These data provide the rational to propose that pro-inflammatory, pro-angiogenic activities by PCa pTA-NKs may also act, by shaping monocyte/macrophage polarization and functions. M2-like macrophages/TAMs have been associated with increased tumour angiogenesis and poorer survival in PCa patients [59–61].

A study showed that IL-2 priming of NK cells from patients with PCa, resulted in distinct NK cell phenotypes and correlates with different NK cytotoxic activities [38]. Once again, these cited results, together with our study, point out the important role of the phenotype and functions of NK cells in PCa patients and other cancer types, that could be used, in the future, as novel cellular circulating biomarker and could be helpful in selecting precise immunotherapeutic approaches and combinations [38].

We extended our investigations using an *in vitro* model, mimicking the interaction of PCa secreted products with normal circulating NK cells. Using three different PCa cell lines, PC-3, DU-145 and LNCaP, respectively, we observed that their conditioned media were able to induce the CD56^bright^CD9^+^CD49a^+^CXCR4^+^ phenotype in NK cells derived from healthy controls. We confirmed that CM from PCa cell lines polarize NK cell, that acquires acquire feature of anergy, with reduced capability to produce the anti-tumour cytokines IFNγ, TNFα and the cytotoxic factor granzyme B. In our model, we also observed that that CM from PCa cell lines induced the dNK-like phenotype on cytolytic NK cells, that are able to produce pro-inflammatory factors, including CXCL8 and CXL12.

In other tumours where pTA-NKs have been investigated, CD9 is present on the cell surface of a subset of circulating CD56^bright^CD16^-/low^; dNK-like cells are present in pNK of melanoma [62–64], lung [12,28,65] and colorectal cancers [25,66]. Based on our evidence we confirmed that, even when it is difficult to detect tumour infiltrating NK, peripheral blood NK cells offer and indirect tool monitor the state of cancer patients according to their polarization.

Here we fill in an existing gap of knowledge regarding pNK polarization in prostate cancer patients as a potential useful biomarker.

## CONCLUSIONS

Our results show that PCa TA-NKs are effector cells able to support inflammation and inflammatory angiogenesis in PCa, by producing factors stimulating endothelial cells, monocytes recruitment and macrophage polarization toward M2 like/TAM phenotype. These data provide a rationale for the future use of pNK pro inflammatory angiogenesis profile in diagnostics and for designing therapies to restore NK lytic activity and polarization in PCa.

## ABBREVIATIONS

ADCC: Antibody Dependent Cellular Cytotoxicity
ADK: Adenocarcinoma
ANG: Angiogenin
ANGIOP1: Angiopoietin 1
ANOVA: Analysis of Variance
CCL: Chemokine Ligand (C-C motif)
cDNA: complementary DNA
CM: Conditioned Media
CRPC: Castration Resistant Prostate Cancer
CXCL: Chemokine Ligand (C-X-C motif)
DNAM-1: DNAX accessory molecule 1
dNK cells: decidual Natural Killer cells
EBM: Endothelial cell Basal Medium
EGM: Endothelial cell Growth Medium
FACS: Fluorescence-Activated Cell Sorting
FBS: Fetal Bovine Serum
FC: Flow Cytometry
FSC: Forward Scatter
GAPDH: Glyceraldehyde-3-Phosphate Dehydrogenase
GM-CSF: Granulocyte-Macrophage Colony-Stimulating Factor
HBV: Hepatitis B Virus
HC: Healthy Control
HCV: Hepatitis C Virus
HIV: Human Immunodeficiency Virus
HUVEC: Human Umbilical Vein Endothelial Cell
ICAM: Inter Cellular Adhesion Molecule-1
IFN-: γ = Interferon γ
IHC: Immuno Histo Chemistry
IL-: Interleukin-
I-TAC: Interferon-inducible T-cell Alpha Chemoattractant
mAbs: monoclonal Antibodies
MCP-1/CCL2: Monocyte Chemoattractant Protein-1
MMPs: Matrix Metallo Proteinases
MNCs: Mono Nuclear Cells
NK: Natural Killer
NKG2D: Natural Killer receptor Group 2 D
NSCLC: Non-Small Cell Lung Cancer
PAI: Plasminogen Activator Inhibitor
PB: Peripheral Blood cells
PBMCs: peripheral Blood Mononuclear Cells
PCa: Prostate Cancer
PD-L1: Programmed Death receptor Ligand 1
PGE2: Prostaglandin E2
PlGF: Placental Growth Factor
PMA: Phorbol Merystate Acetate
P/S: Penicillin / Streptomycin
PTA-NKs: Prostate Tumor Associated Natural Killer cells
RANTES: Regulated upon Activation, Normal T Cell Expressed and Presumably Secreted
SSC: Side Scatter
STAT: Signal Transducer and Activator of Transcription
TAMs: Tumor Associated Macrophages
TGFβ: Transforming Growth Factor-β
TIMP: Tissue Inhibitor of Metallo-Proteinase
TME: Tumor Microenvironment
TNF α: Tumor Necrosis Factor-α
Treg: T Regulatory Cells
uPAR: urokinase-type Plasminogen Activator Receptor
VCAM: Vascular Cellular Adhesion Molecule-1
VEGF: Vascular Endothelial Growth Factor
VEGFR: Vascular Endothelial Growth Factor Receptor

## AKNOWLEDGEMENTS

We thank Dr. Paola Corradino, IRCCS MultiMedica, Milan, Italy, for support to literature research. We thank Dr. Barbara Basani, IRCCS National Cancer Institute, Milan, Italy, for helpful discussion and revision of the manuscript.

## AUTHOR CONTRIBUTIONS

DB, MG, LM, ABo, GB, AB: performed the experiments. DB, AB, analysed data, perform the statistical analysis, prepared figures. FD, PC, AN, AG: provided and collected the clinical samples and provide the clinical support. DB, LM, DMN, AA, AB: conceived the experiments, analysed data. DB, LM, DMN, AA, AB: wrote the manuscript. DMN, AB: provided funds.

---

**Supplementary Figure 1. Expression of CD16 and NKG2D in PCa pTA-NKs and controls.** TANKs from PCa pNK patients share similar pNK cell subset frequency (CD56^+^CD16^+^ and CD56^+^CD16^-^ cells) in peripheral blood samples (A). The NKG2D activation marker is downregulated in PCa ADK pTA-NKS as compared with those from healthy controls (B). Data are showed as mean ± SEM, t-test student, **p<0.01. HC: healthy controls; ADK: prostate adenocarcinoma

**Supplementary Figure 2. Overall secretome array**. Scan acquisition for the angiogenesis antibody-array (C1 and C2), showing overall dots, following exposure to secreted products (CM) from FACS sorted pNK cells of PCa-ADK patients and controls. Tables to identify dots position for array C1 and C2 is provided. CMs were pooled from pNK cells sorted from 3 different PCa patients. Arrays were performed in duplicates. ADK: prostate adenocarcinoma; HC: healthy controls.

**Supplementary Table1. Demographic features of our cohort of PCa patients and controls**. Table summarizing the features of our cohorts of patients, with relative sample size. Average of age is showed as mean ± SD. N: sample size, ADK: prostate adenocarcinoma; HC: healthy controls.

**Supplementary Table 2. Primer sequences for oligos used for Real-time PCR**. Sequences for forward and reverse oligos used for real-time PCR are showed.

**Supplementary Table 3. Antibodies used in flow cytometry experiments.** The table summarize the antibodies (primary conjugated, primary not-conjugated, secondary conjugated) used in flow cytometry analysis.

