## Supplementary material for "Prostate cancer peripheral blood NK cells show enhanced CD9, CD49a, CXCR4, CXCL8, MMP-9 production, and secrete monocyte-recruiting and polarizing factors": Supplemetary Material

Supplementary Figure 1

A

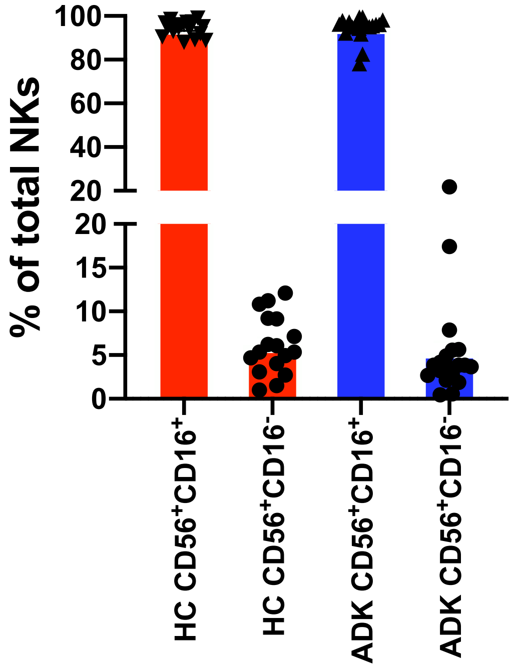


B

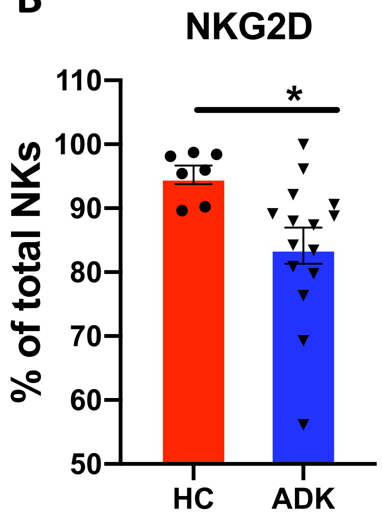

### Supplementary Figure 2

**ARRAY C1**

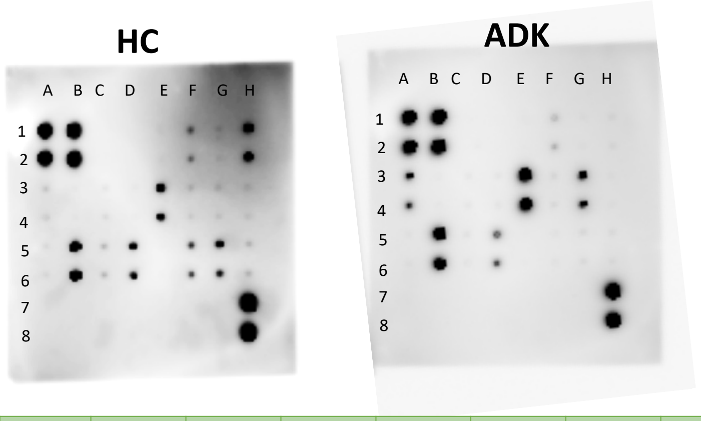


| | A | B | C | D | E | F | G | H |
|---|---|---|---|---|---|---|---|---|
| 1<br>2 | POS | POS | NEG | NEG | ANG | EGF | CXCL5 | bFGF |
| 3<br>4 | GRO | IFNγ | IGF-1 | IL-6 | IL-8 | LEPTIN | CCL2 | PDGF-BB |
| 5<br>6 | PIGF | CCL5 | TGFβ | TIMP-1 | TIMP-2 | TPO | VEGF | VEGF-D |
| 7<br>8 | BLANK | BLANK | BLANK | BLANK | BLANK | BLANK | NEG | POS |

**ARRAY C2**

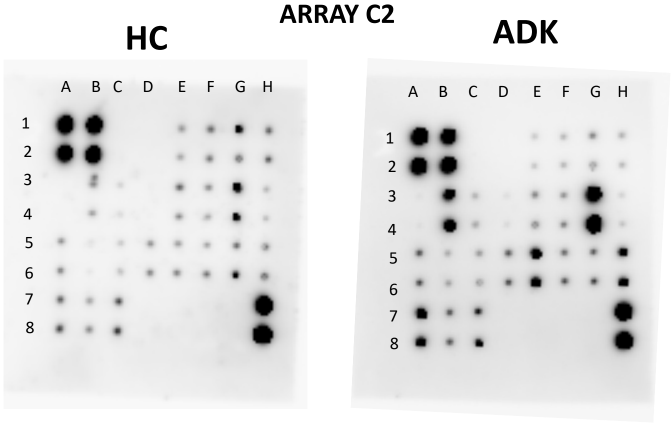


| | A | B | C | D | E | F | G | H |
|---|---|---|---|---|---|---|---|---|
| 1<br>2 | POS | POS | NEG | NEG | ANGPT-1 | ANGPT-2 | PLG | ENDOST |
| 3<br>4 | G-CSF | GM-CSF | CCL1 | IL-10 | IL-1α | IL-1β | IL-2 | IL-4 |
| 5<br>6 | CXCL11 | CCL7 | CCL13 | MMP-1 | MMP-9 | CD31 | TIE-2 | TNFα |
| 7<br>8 | uPAR | VEGF R2 | VEGF R3 | BLANK | BLANK | BLANK | BLANK | POS |

#### Supplementary Table 1

| DIAGNOSIS | ADK | HC (Ctrl) |
|---|---|---|
| N° of subjects | 25 | 20 |
| Age (mean ± sd) | 69±8.36 | 57±8.36 |

#### Supplementary Table 2

| TABLE S1: human primers for qRT-PCT | |
|---|---|
| ***Gene*** | ***Sequences*** |
| **hu 18s** | forward: *gcagaatccacgccagtacaag* |
| | reverse: *gcttgttgtccagaccattggc* |
| **hu plgf** | forward: *tgtcaccatgcagctcctaa* |
| | reverse: *gtctgtgggtctctgcttct* |
| **hu vegf** | forward: *ctgtcttgggtgcattggag* |
| | reverse: *accagggtctcgattggatg* |
| **hu cxcl8** | forward: *cctgatttctgcagctctgtg* |
| | reverse: *gtggtccactctcaatcactctc* |
| **hu vegfr2** | forward: *tctctctgcctacctcacct* |
| | reverse:*accataccactgtccgtctg* |
| **hu cxcl12** | forward: *ctcaacactccaaactgtgccc* |
| | reverse: *ctccaggtactcctgaatccac* |
| **hu pai** | forward: *tctctgccctcaccaacatt* |
| | reverse: *cggtcattcccaggttctct* |
| **hu icam-1** | forward: *agcggctgacgtgtgcagtaat* |
| | reverse: *tctgagacctctggcttcgtca* |
| **hu vicam-1** | forward: *gattctgtgcccacagtaaggc* |
| | reverse: *tggtcacagagccaccttcttg* |
| **hu cxcr4** | forward: *ctcctctttgtcatcacgcttcc* |
| | reverse: *ggatgaggacactgctgtagag* |
| **hu il-1α** | forward: *tgtatgtgactgcccaagatgaag* |
| | reverse: *agaggaggttggtctcactacc* |
| **hu arg1** | forward: *gattctcagtgctgcggatc* |
| | reverse: *cagcttctcttatggcagcg* |
| **hu il10** | forward: *gccaagccttgtctgagatg* |
| | reverse: *aagaaatcgatgacagcgcc* |
| **hu il12** | forward: *gccttcaccactcccaaaac* |
| | reverse*: atggtaaacaggcctccact* |
| **hu cd206** | forward: *agccaacaccagctcctcaaga* |
| | reverse: *caaaacgctcgcgcattgtcca* |

**Supplementary table 3**

| MAB | CLONE | REACTIVITY | FLUOROCROME | TARGET | SUPPLIER |
|---|---|---|---|---|---|
| CD184/CXCR4 | 12GS | Human | PE | C-X-C chemokine receptor type 4 | Miltenyi Biotec |
| CD314/NKG2D | REA122 8 | Human | PE | | Miltenyi Biotec |
| Ang | 14017.7 | Human | _ | Angiogenin | Abcam |
| Angiop1 | | Human | _ | Angiopoietin-1 | Abcam |
| CD16/FcγRIII | REA423 | Human | FITC | Fc-gamma-ReceptorIII | Miltenyi Biotec |
| CD3 | BW264/5 6 | Human | PerCP | T-cell receptor (TCR) | Miltenyi Biotec |
| CD49a | REA110 6 | Human | PE | α1 integrin | Miltenyi Biotec |
| CD56 | REA196 | Human | APC | Neural cell adhesion molecule (NCAM | Miltenyi Biotec |
| CD9 | REA107 1 | Human | PE | Tetraspanin | Miltenyi Biotec |
| CXCL12/SDF-1 | 79018 | Human | PE | C-X-C Motif Chemokine Ligand 12/Stromal cell-derived factor-1 | R&D SYSTEM |
| CXCL8/IL8 | E8N1 | Human | PE | C-X-C Motif Chemokine Ligand 8/Interleukin-8 | Miltenyi Biotec |
| GRZ-B | CB9 | Human | PE | Granzyme-A | Miltenyi Biotec |
| IFNγ | 4S.B3 | Human | PE | Interferon-gamma | Miltenyi Biotec |
| TNFα | Mab11 | Human | PE | Tumor Necrosis Factor-alpha | Miltenyi Biotec |
| VEGF | # 23410 | Human | PE | Vascular Endothelial Growth Factor | R&D SYSTEM |
